# Let’s Tweap again: Economic and SNP retrieval optimisation for target enrichment of ancient DNA

**DOI:** 10.64898/2026.01.15.699081

**Authors:** Tina Saupe, Mariam Omar Gómez, E-Jean Tan, Helja Kabral, Christiana L. Scheib, Carolina Bernhardsson, Mattias Jakobsson

## Abstract

Advancements in ancient DNA (aDNA) research have enabled the recovery of individual human genomes through target enrichment approaches. Despite the decreasing cost of next-generation sequencing, enrichment methods targeting single-nucleotide polymorphisms (SNPs) compiled in RNA/DNA hybrid panels remain costly and require further authentic optimisations.

Here, we present the Tweap (<u>Tw</u>ist ch<u>eap</u>) protocol as a cost-effective alternative to the commercialised Twist protocol for target enrichment via in-solution hybridisation of ancient human DNA by replacing the streptavidin-coated superparamagnetic beads for binding the hybridised targeted DNA. We evaluated the protocol on 14 double-stranded DNA libraries from ancient human individuals originating from a hot and humid climate zone known to accelerate DNA degradation. The selected individuals reflected similar preservation levels and a broad range of endogenous human DNA content (0.79 – 41.69 %).

We optimised and simplified both the Twist and Tweap protocols for target enrichment, particularly for libraries with ultra-low library complexity, making it reproducible in any life sciences laboratory. The modified Tweap protocol not only reduced overall reagent costs and consumption, contributing to more climate-conscious research, but also recovered up to 7% more unique SNPs when applying multi-pooling of libraries from the same individual.

Our findings demonstrate that the Tweap protocol is a viable, efficient, and scalable alternative for SNP capture in ancient DNA (aDNA) studies, supporting broader accessibility and sustainability in ancient DNA research.

## Introduction

Many ancient DNA (aDNA) studies focus on (partial) genome recovery of skeletal remains from various geographical regions in one or multiple climatic zones. Until today, the recovery of genomes from areas closer to the equator, where climatic conditions are hot and humid, has been challenging due to poor preservation of the skeletal remains and rapid DNA damage and degradation rates (Cuesta-Aguirre *et al*., 2025).

Therefore, countless studies have been published from the Northern Hemisphere, leading to profound gaps in the reconstruction of the human past (Mallick *et al*., 2024). Nevertheless, with the increasing interest in these challenging regions, new laboratory methods have been developed to recover the DNA from ancient individuals. One of the more recently used methodologies in aDNA research is applying a target enrichment method that allows researchers to target DNA fragments of interest using panels containing biotinylated single-stranded (ss) or double-stranded (ds) DNA, or RNA sequences (Gnirke *et al*., 2009).

In the last years, researchers have increasingly applied biotinylated dsDNA probes from Twist Bioscience (San Francisco, USA) to enrich aDNA libraries with low endogenous human DNA content (endo%), thereby reducing the allelic bias seen in previously available probe panels (Davidson *et al*., 2023, 2024). A customised ‘Ancient Human DNA’ panel (hereafter: ‘1350k’) was developed by Twist Bioscience to reduce the allelic bias towards enriching one allele over another, leading to higher coverage and uniformity. The panel targets 1,352,535 SNPs merged into 1,434,155 probes. Twist Bioscience also offers an additional ‘mitochondrial DNA’ (mtDNA) panel covering all 16,659 base pairs (bp) and 37 genes. The efficiency of the ‘1350k’ panel has been compared to two other panels: the ‘1240k’ panel, which cannot be purchased, and the Daicel Arbor Biosciences ‘myBaits Expert Human Affinities Prime Plus’ reagent, which is no longer available (Rohland *et al*., 2022a). The authors concluded that the capture performance of the ‘1350k’ panel was more proficient for uracil DNA glycosylase (UDG) treated ssDNA and partial-UDG-treated dsDNA libraries and that their DNA libraries with low endogenous human DNA content required only one round of hybridisation reaction instead of two (Carpenter *et al*., 2013).

In a thorough assessment of the modified ‘Twist Target Enrichment Standard Hybridization v1’ protocol optimised for aDNA libraries (Rohland *et al*., 2022a), we noticed that this protocol required three times more streptavidin-coated binding beads (part of the ‘Twist Dry Down Beads’ kit, #104325) and binding buffer (part of the ‘Twist Hybridisation and Wash Buffer’ kit, #101025) to capture the targets after the hybridisation step (Figure 1). On the contrary, only 1 µl of the ‘Human Ancient DNA’ panel (instead of the 4 µl required for modern DNA libraries) and 0.167 µl of the ‘mtDNA’ panel are required for ss/dsDNA libraries of ancient individuals, allowing researchers to potentially perform four times more hybridisation reactions. However, a 12- or 96-reaction Twist Bioscience kit including the streptavidin binding beads and binding buffers would only allow four or 24 hybridisation reactions, respectively.

**Figure 1.**
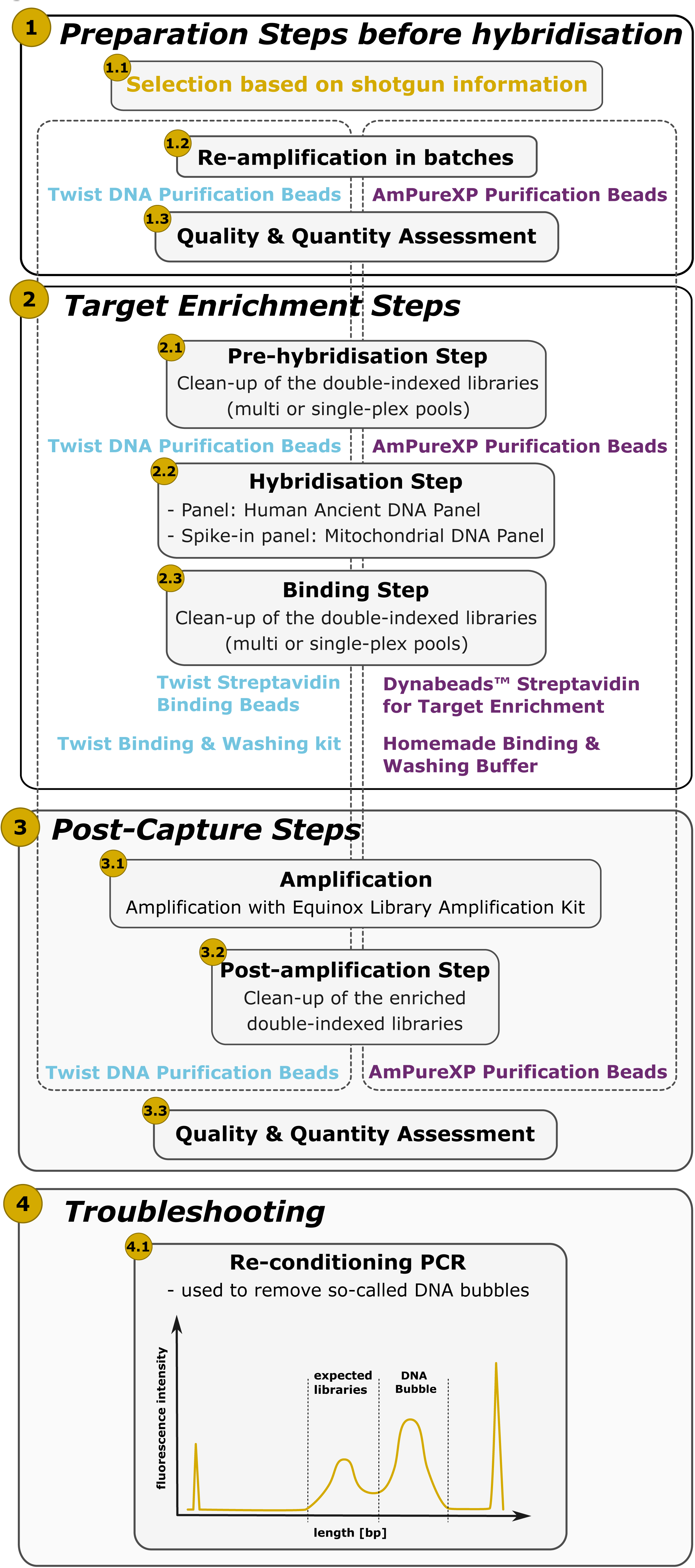
Workflow target enrichment via in-solution hybridisation using Twist or Tweap protocols, and their differences are marked in cyan and purple text, respectively. More information is in Supplementary Material 1.

Therefore, using the entire "Human Ancient DNA" panel to enrich additional aDNA libraries would require tripling the purchase of streptavidin binding beads and binding buffers (Table 1).

**Table 1.** Overview of required reagents to perform a target enrichment via in-solution hybridisation using the Twist ‘Ancient Human DNA’ panel for 12 hybridisation reactions. *The customised ‘Ancient Human DNA’ Panel was only available as a 96-hybridisation reaction kit in 2023. *rxn -* reaction.

|  | Tweap Protocol | Twist Protocol |
| --- | --- | --- |
| **Probe panels** |  |  |
| ‘Ancient Human DNA’ Panel* | 1 | 1 |
| ‘Mitochondrial DNA’ Panel (12 rxn) | 1 | 1 |
| **Hybridisation step reagents** |  |  |
| Twist Universal Blockers (12 rxn) | 1 | 1 |
| Twist Hybridisation Reagents (12 rxn) | 1 | 1 |
| **Binding & Washing and post-amplification steps reagents** |  |  |
| Twist Dry Down Beads kit (12 rxn) (including Streptavidin binding beads and DNA Purification beads) | not required | 3 |
| Twist Wash Buffers kit (12 rxn) | not required | 3 |
| Dynabeads for target enrichment (2 ml) | 1 | not required |
| AmPure XP beads (60 ml) | 1 | not required |

Here, we adapted the Twist protocol (standard v1 using the Rohland et. al 2022 modification) to enrich dsDNA libraries with ultra-low endogenous human DNA content (as low as 0.79 %) by replacing the DNA purification and streptavidin binding beads with comparable materials from alternative suppliers (See Supplementary Material 1). This allowed us to cost-efficiently increase the number of hybridisation reactions using the ‘Human Ancient DNA’ panel, reducing reagent costs (up to 45 %) and chemical waste in life science laboratories (Figure 1). We examined published protocols using streptavidin binding beads for targeting biotinylated probes bound to DNA, adapted by many researchers working with low DNA content materials (Furtwängler et al., 2020; Carpenter et al., 2013; González Fortes & Paijmans, 2019; Horn, 2012; Shapiro & Hofreiter,(Horn, 2012; Shapiro and Hofreiter, 2012; Carpenter *et al*., 2013; González Fortes and Paijmans, 2019; Furtwängler *et al*., 2020). For example, in the ‘Ancient DNA - Methods and Protocols’ book, various Dynabeads products from Invitrogen, such as the M-270 Streptavidin and MyOne Streptavidin C1 beads, were described to enrich ancient dsDNA libraries using different probe panels (Shapiro and Hofreiter, 2012). Based on the efficiency rates of the Dynabeads described in the protocols above, we selected the Dynabeads Streptavidin for Target Enrichment beads, presenting a high binding capacity (up to 20 µg of biotinylated dsDNA), to replace the Twist streptavidin binding beads. This crucial change eliminated the whole requirement to purchase the ‘Twist Dry Down Beads’ kit, which includes the Streptavidin binding beads and DNA Purification beads (Table 1). Following this, we obligatorily substituted the Twist DNA purification beads with AmPure XP beads, which are commonly used in our facility, but not exclusively required for the efficiency of the protocol presented here, and therefore replaceable with any DNA purification beads of choice. In this study, we investigated the efficiency of our newly refined <u>Tw</u>ist ch<u>eap</u> (Tweap) and the Twist protocols by applying both protocols to an experimental set of dsDNA libraries from previously unpublished ancient human individuals with poor DNA preservation, dated to around 3,500 years ago, with full consent from the stakeholders of the individuals.

In the present publication, we focus on optimising the protocol(s) to enhance the quality of the retrieved genome-wide SNP data. Any archaeological contextual information and generated sequencing data will be provided in future publications, where the focus will be on the analysis of their population history. In addition to the reagent replacements that resulted in reduced costs (approximately 45%), we focused on sustainable consumption of reagents and minimising waste in the laboratory workflow (Table 1).

## Methods

### Selection criteria for the target enrichment via in-solution hybridisation

To validate the selection criteria for sufficient target enrichment, we generated an experimental set of blunt-end, double-indexed dsDNA libraries from ancient human remains at the Ancient DNA facility at Uppsala University (Sweden), using DNA extracts generated at the Ancient DNA facility, Institute of Genomics, University of Tartu (Estonia). These were sequenced on a next-generation sequencing (NGS) platform (Illumina NovaSeq 6000 Sequencing system or NovaSeq X Plus Sequencing system) at SNP&SEQ Technology Platform, Uppsala University (Sweden) before the target enrichment via in-solution hybridisation was performed (See Supplementary Material 1 for details, Supplementary Data SD1A-C).

For the validation, we focused on the statistical analysis of the sequenced reads obtained from the shotgun sequenced dsDNA libraries and aligned to the reference human genome build (hg19/hs37d5) using an in-house script exploring the percentages of human mapping proportion/endogenous human DNA content [%] (*mapped reads, including duplicates and short reads, divided by trimmed reads)* and the library complexity [%] (*final/non-duplicated mapped reads divided by trimmed reads)*, as well as the short read (< 35 base pairs (bp)) and PCR duplicate rates [%] and the ratio between mtDNA: nuclear DNA reads (Table 2, Supplementary Data SD1D) (Hernandez-Rodriguez *et al*., 2018). First, we evaluated the human mapping proportion and the library complexity of each dsDNA library, taking into account that researchers use the outputs after deduplication for further downstream analysis. In this regard, a high human mapping proportion may suggest efficient target enrichment; however, this ratio typically includes the PCR duplicates and short reads, which could be high and therefore result in a lower library complexity.

**Table 2.** Detailed information on selecting dsDNA libraries for target enrichment via in-solution hybridisation. *Conc -* concentration; *n -* number.

| Criteria | Definition | Validation/Recommendation |
| --- | --- | --- |
| **Obtained from the shotgun sequencing information.** |  |  |
| Average length distribution | A typical aDNA library has an average length of 40-100 bp (Rasmussen et al. 2010). | Unusual average length - check for present-day human contamination. |
| Deamination profile/rate | The deamination profile presents increased C→T substitutions at the 5'-end and G→A substitutions at the 3'-end. The average deamination rates are calculated using the first 25 bp of the DNA reads (Jónsson et al. 2013; Ginolhac et al. 2011). | For blunt-end repaired dsDNA libraries of ancient individuals, the deamination rates can vary between 10 % and 50 %, presented in a smooth curve. Libraries with few DNA reads also displayed similar curves, but with background noise. |
| Human proportion [%] | The protion of human DNA including PCR duplicates and short reads mapped to the human reference genome. | The recommendation for sufficient enrichment is >1%, however, if the individual is of important/great interest, an attempt at enrichment could be beneficial. |
|  | Calculation:<br>library complexity/unique DNA fraction = unique n of reads/n of aligned reads |  |
| library complexity/unique DNA fraction [%] | The portion of unique human DNA in a dsDNA library reflects an ancient individual's overall human DNA preservation. Value | ↓ value = low amount of unique DNA molecules available for deeper sequencing or target enrichment |
|  | Calculation:<br>library complexity/unique DNA fraction = unique n of reads/n of aligned reads | ↑ value = high amount of unique/diverse DNA molecules available for deeper sequencing and target enrichment |
| mtDNA/nuclear DNA ratio | Calculated based on the number of mapped unique reads to the nuclear and mitochondrial genomes. | ↓ rratio = more nuclear than mitochondrial DNA sequences available |
|  |  | ↑ rratio = more mitochondrial than nuclear DNA sequences available |
|  | Calculation:<br>Rratio = Rmitochondrial/Rnuclear |  |
| Number of single nucleotide polymorphisms | Estimation of the number of SNPs using the respective SNP panel of interest. | If the number of SNPs is low or none, the capture may not be able to enrich efficiently for genome-wide data analysis. |
| PCR duplicate rate [%] | PCR duplicate rate is calculated using the mapped reads to the reference human genome with identical start and end positions. Researchers often refer to it as ‘clonality’. | A high PCR duplication rate suggests an over-amplification of the dsDNA library, which would increase during enrichment/deep sequencing, leading to increased costs. |
| Present-day human mtDNA contamination estimation (if applicable) |  | <5% = pass quality filter |
|  |  | 5-10% = acceptable contamination, but careful interpretation is needed |
|  |  | >10% = additional filter required or excluded from genome-wide data analysis |
| Obtained from the quality & quantity assessment of the dsDNA libraries. |  |  |
| DNA concentration [ng/μl] | The DNA concentration is the total amount of DNA in a dsDNA library after amplification and clean-up, usually measured using a Qubit® Fluorometer. The value does not exclude adapter dimers, amplification artefacts, and deficient clean-up performance. | DNA concentration can be high, even if the endogenous DNA content is (ultra) low. High DNA concentration minimises over-amplification of off-target DNA and more favourable DNA fragments for amplification. |
| Fragment analysis using e.g. TapeStation | A good quality dsDNA library usually presents a nice peak around 180-220 bp and no adapter dimers or amplification artefacts. | If the fragment analysis presents a secondary peak around 400 - 1000 bp, a 'bubble DNA' phenomenon, a re-conditioning PCR step can be performed to reduce this (See Supplementary Material 1). |
| Target DNA amount [ng] | The target DNA amount is based on the DNA concentration and the minimum volume of 10 μl. Following the Twist recommendation, 500 - 1000 ng of DNA is required for an efficient enrichment.<br>Calculation:<br>Target DNA conc = DNA conc *input volume [μl] | If the amount of DNA is low, it is recommended to apply multiple re-amplifications of the dsDNA libraries to increase the target DNA, while the 'Alternate pre-hybridisation DNA concentration' protocol demonstrates efficient enrichment up to 40 μl starting volume. |

In addition, an initial high percentage of PCR duplicates may reflect the possible over-amplification of PCR duplicates during the re-amplification and post-capture amplification steps (Tables 2 and 3, Supplementary Material 1, Supplementary Data SD1). While shorter reads during the hybridisation reactions are less likely to be enriched, the number is relevant to pre-evaluate the efficiency of sufficient target enrichment. Therefore, enriching a library with a high human mapping proportion might be efficient; however, if the library complexity is low, the enrichment may result in fewer unique SNPs than expected. In addition, we examined the raw number of SNPs and the read-based SNP discovery (*number of SNPs divided by number of targeted reads*) on the ‘1350k’ panel to determine whether a dsDNA library is likely to capture an adequate number of SNPs for panel-restricted analysis (Supplementary Data SD1E-F). In summary, the experimental set consisted of aDNA libraries of 14 unique human individuals with various human proportions ranging between 0.79 and 41.69 % (median = 6.01 %) and library complexities between 0.09 and 30.74 % (median = 2.32 %) in addition to the low modern human contamination on the mitochondrial genome (0.86 – 14.48 %, median = 4.78 %) (Table 3, Supplementary Data SD1). All aDNA libraries exhibited deamination patterns (mean [C>T] = 26.47%, mean [G>A] = 26.37%), which are characteristic of ancient DNA (Supplementary Figure S1) (Ginolhac *et al*., 2011a; Jónsson *et al*., 2013a).

**Table 3.**
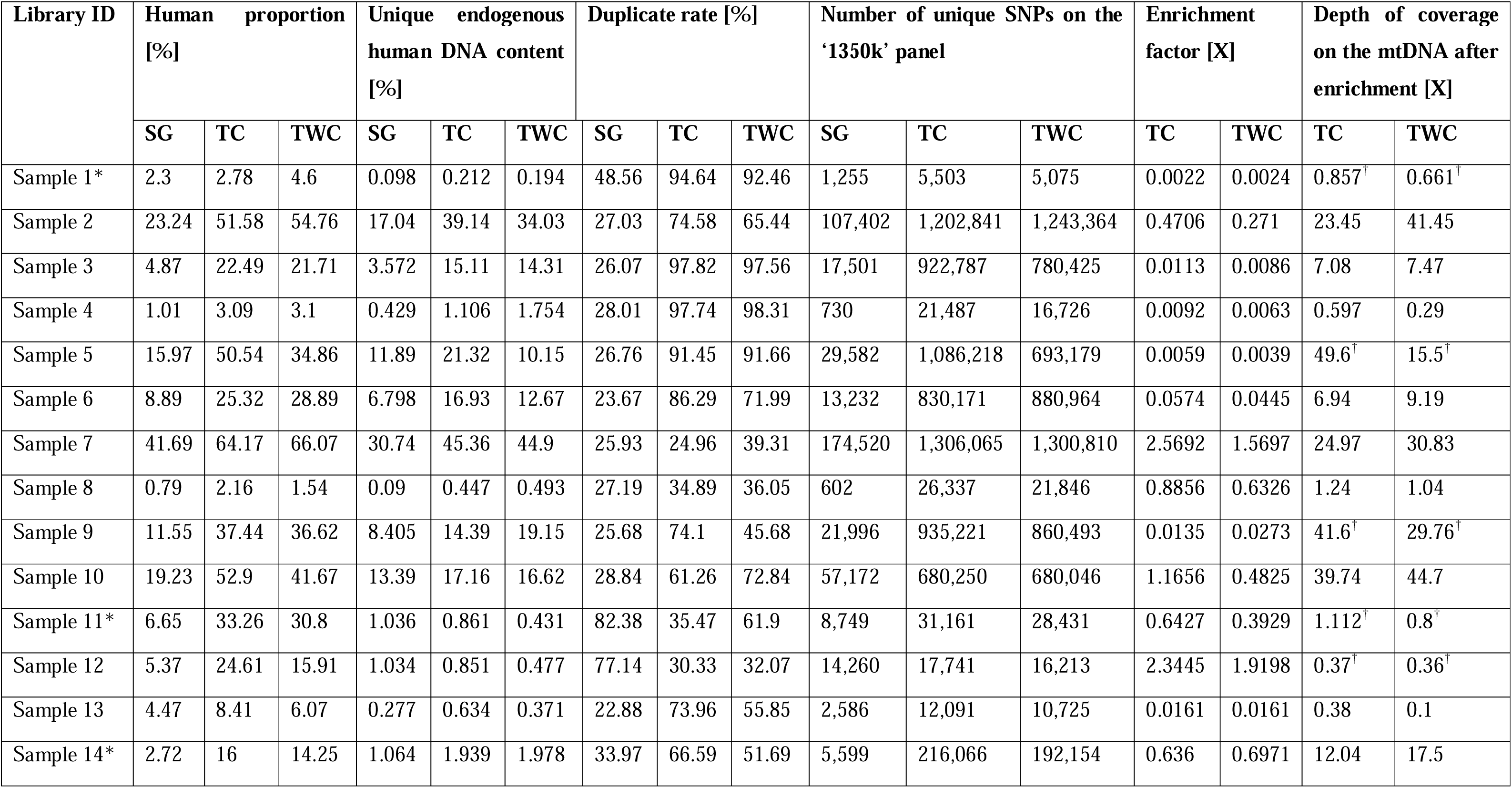
Statistical information of the selected samples before and after the enrichment included in this study (Supplementary Material 1, Supplementary Data SD1). Samples with ‘*’ are multi-pool hybridisation reactions. *SG* - Shotgun sequencing; *SNPs* - Single nucleotide polymorphisms; *TC* - Twist capture; *TWC* - Tweap capture. ^†^ = spike-in of 0.5 µl ‘mtDNA’ panel.

| Library ID | Human proportion [%] |  |  | Unique endogenous human DNA content [%] |  |  | Duplicate rate [%] |  |  | Number of unique SNPs on the ‘1350k’ panel |  |  | Enrichment factor [X] |  | Depth of coverage on the mtDNA after enrichment [X] |  |
| --- | --- | --- | --- | --- | --- | --- | --- | --- | --- | --- | --- | --- | --- | --- | --- | --- |
|  | SG | TC | TWC | SG | TC | TWC | SG | TC | TWC | SG | TC | TWC | TC | TWC | TC | TWC |
| Sample 1* | 2.3 | 2.78 | 4.6 | 0.098 | 0.212 | 0.194 | 48.56 | 94.64 | 92.46 | 1,255 | 5,503 | 5,075 | 0.0022 | 0.0024 | 0.857<sup>†</sup> | 0.661<sup>†</sup> |
| Sample 2 | 23.24 | 51.58 | 54.76 | 17.04 | 39.14 | 34.03 | 27.03 | 74.58 | 65.44 | 107,402 | 1,202,841 | 1,243,364 | 0.4706 | 0.271 | 23.45 | 41.45 |
| Sample 3 | 4.87 | 22.49 | 21.71 | 3.572 | 15.11 | 14.31 | 26.07 | 97.82 | 97.56 | 17,501 | 922,787 | 780,425 | 0.0113 | 0.0086 | 7.08 | 7.47 |
| Sample 4 | 1.01 | 3.09 | 3.1 | 0.429 | 1.106 | 1.754 | 28.01 | 97.74 | 98.31 | 730 | 21,487 | 16,726 | 0.0092 | 0.0063 | 0.597 | 0.29 |
| Sample 5 | 15.97 | 50.54 | 34.86 | 11.89 | 21.32 | 10.15 | 26.76 | 91.45 | 91.66 | 29,582 | 1,086,218 | 693,179 | 0.0059 | 0.0039 | 49.6<sup>†</sup> | 15.5<sup>†</sup> |
| Sample 6 | 8.89 | 25.32 | 28.89 | 6.798 | 16.93 | 12.67 | 23.67 | 86.29 | 71.99 | 13,232 | 830,171 | 880,964 | 0.0574 | 0.0445 | 6.94 | 9.19 |
| Sample 7 | 41.69 | 64.17 | 66.07 | 30.74 | 45.36 | 44.9 | 25.93 | 24.96 | 39.31 | 174,520 | 1,306,065 | 1,300,810 | 2.5692 | 1.5697 | 24.97 | 30.83 |
| Sample 8 | 0.79 | 2.16 | 1.54 | 0.09 | 0.447 | 0.493 | 27.19 | 34.89 | 36.05 | 602 | 26,337 | 21,846 | 0.8856 | 0.6326 | 1.24 | 1.04 |
| Sample 9 | 11.55 | 37.44 | 36.62 | 8.405 | 14.39 | 19.15 | 25.68 | 74.1 | 45.68 | 21,996 | 935,221 | 860,493 | 0.0135 | 0.0273 | 41.6<sup>†</sup> | 29.76<sup>†</sup> |
| Sample 10 | 19.23 | 52.9 | 41.67 | 13.39 | 17.16 | 16.62 | 28.84 | 61.26 | 72.84 | 57,172 | 680,250 | 680,046 | 1.1656 | 0.4825 | 39.74 | 44.7 |
| Sample 11* | 6.65 | 33.26 | 30.8 | 1.036 | 0.861 | 0.431 | 82.38 | 35.47 | 61.9 | 8,749 | 31,161 | 28,431 | 0.6427 | 0.3929 | 1.112<sup>†</sup> | 0.8<sup>†</sup> |
| Sample 12 | 5.37 | 24.61 | 15.91 | 1.034 | 0.851 | 0.477 | 77.14 | 30.33 | 32.07 | 14,260 | 17,741 | 16,213 | 2.3445 | 1.9198 | 0.37<sup>†</sup> | 0.36<sup>†</sup> |
| Sample 13 | 4.47 | 8.41 | 6.07 | 0.277 | 0.634 | 0.371 | 22.88 | 73.96 | 55.85 | 2,586 | 12,091 | 10,725 | 0.0161 | 0.0161 | 0.38 | 0.1 |
| Sample 14* | 2.72 | 16 | 14.25 | 1.064 | 1.939 | 1.978 | 33.97 | 66.59 | 51.69 | 5,599 | 216,066 | 192,154 | 0.636 | 0.6971 | 12.04 | 17.5 |

### The implementation of the protocols

To test the performance of the two methods, we applied the modified published protocol ‘Twist Target Enrichment Standard Hybridization v1’ (https://www.twistbioscience.com/resources/protocol/twist-target-enrichment-standard-hybridization-v1-protocol; Revision 4.0) and our Tweap protocol on the selected dsDNA libraries in the post-PCR laboratory at the Ancient DNA facility, Uppsala University (Sweden) (Figure 1, See Supplementary Material 1 for details) (Rohland *et al*., 2022a). A step-by-step format of the Tweap protocol used in this article is also available on protocols.io (https://www.protocols.io/blind/2107D8C61E8E11F0A7670A58A9FEAC02). We conducted 11 single-pool (identical to single-plex) and three multi-pool (samples: 1, 11, and 14) hybridisation reactions using the same input DNA concentrations reflected in the starting volume when performing both protocols (Supplementary Figure S2). We multi-pooled dsDNA libraries of unique samples with ∼1% of endogenous human DNA content (Supplementary Material 1, Supplementary Data SD1). To enrich the dsDNA libraries in multi-pool hybridisation reactions equally well, four PCR amplifications built from two dsDNA libraries of the same individuals were pooled in equal molar amounts before the start of the target enrichment (See Supplementary Material 1 for details). The Twist protocol recommends an input concentration between 500 and 1000 ng DNA to ensure a sufficient hybridisation reaction for modern samples. However, ancient DNA libraries are often scarce, and their measured DNA concentration, typically very low, reflects a mixture of various DNA sources, such as from the environment, bacteria/viruses, and fungi, and not exclusively human DNA (Marciniak *et al*., 2015). To compensate for the presence of low human DNA molecules, a re-amplification step was performed to increase the amount of DNA fragments of interest and improve both the target enrichment efficiency and library complexity before enrichment (See Supplementary Material 1). To prevent the over-amplification of shorter fragments, GC-rich fragments and/or high-abundance initial fragments, we performed multiple re-amplifications (2 – 4) for one dsDNA library (input: 5 – 10 µl) to maximise the efficiency of the DNA polymerase and decrease the number of low-frequency DNA targets. This step also reduces high duplication rates in samples with low DNA yields. The re-amplified libraries were pooled together if the amplification was sufficient, and a total of 1000 ng was attained. Due to the lower quantities of targeted human DNA fragments in ultra-low endogenous human DNA libraries, over-amplification of such libraries was avoided, using no more than 5 PCR cycles (See Supplementary Material 1). We used different starting volumes (10 – 40 µl) to ensure a minimum amount of 500 ng per selected dsNDA library, applying the ‘Alternate pre-hybridisation DNA concentration’ protocol appended to the Twist Bioscience protocol to concentrate and elute the DNA in 10 µl blocking solution master mix (See Supplementary Material 1, Supplementary Data SD1).

During the quality and quantity assessment of the dsDNA libraries after re-amplification and post-enrichment amplification, we found many samples with elevated DNA concentrations presenting a so-called ‘bubble DNA’ artefact using a fragment analyser. To minimise the presence of such secondary artefact peaks, we applied re-conditioning PCR on dsDNA libraries when the secondary peak was bigger than the primary peak (See Supplementary Material 1 for details) (Thompson, Marcelino and Polz, 2002a).

After the target enrichment, we pooled the enriched dsDNA libraries in equimolar quantities. In the case of multi-pool enriched dsDNA libraries, we considered the DNA concentrations between the single dsDNA libraries after the enrichment to be similar (Supplementary Figure S2). The pools were sequenced on an Illumina NovaSeq X 10B at the SNP&SEQ Technology Platform, Uppsala University (Sweden). We sequenced 46 enrichments from 14 enriched dsDNA libraries (single-pool = 22, multi-pool = 24) (See Supplementary Material 1).

## Results

### Performance of the target enrichment

We evaluated the efficiency of the target enrichment protocols using the quality and quantity assessment of the dsDNA libraries pre- and post-enrichment and the sequencing information after alignment to the human reference genome (See Supplementary Material 1, Supplementary Data SD1). The DNA concentrations of the selected dsDNA libraries before the enrichment ranged between 5.12 and 62.6 ng/µl (median = 23.30 ng/µl). We used between ∼350 and 980 ng of DNA material (median = 701 ng) for the enrichment. Due to the limited input volume available, we did not have the minimum amount of 500 ng DNA for four selected dsDNA libraries (samples 2, 5, 7, and 13); however, the endogenous human DNA content of those (excluding sample 13) was superior to the other samples, indicating that the enrichment might still be successful. After the enrichment, the DNA concentrations ranged between 0.556 and 55.4 ng/µl (median [total] = 17.75 ng/µl, median [Tweap] = 6.38 ng/µl, median [Twist] = 25.6 ng/µl), indicating that the human DNA in the libraries were possibly enriched and amplified (Supplementary Data SD1).

However, the DNA concentrations of the enriched dsDNA libraries with low endogenous human DNA content did not increase significantly, even though the input amount of DNA was close to the total DNA amount recommended by Twist. Thus, we wanted to evaluate the previous theory that the enrichment efficiency depends on the availability of DNA templates using a linear regression model (human mapping proportion [pre-enrichment] ∼ amount of DNA [ng] and library complexity [pre-enrichment] ∼ amount of DNA [ng]) (Cruz-Dávalos *et al*., 2017) (Supplemental Material 1 for details). The results showed that there is a statistically significant negative relationship between the total amount of DNA and the human mapping proportion (adjusted R-squared: 0.1292, p-value = 0.006986) as well as the library complexity (adjusted R-squared: 0.1551, p-value = 0.02178). Those tests suggest that dsDNA libraries with low human DNA templates were most likely highly diluted in relation to the exogenous DNA content, which potentially affected the enrichment efficiency. Furthermore, the over-amplification to reach 1000 ng of input DNA resulted in increased PCR duplicates of both human and non-human DNA, especially in cases where the latter was more prone to amplification. We further tested the relationships between the human proportion (adjusted R-squared = 0.8117, p-value = 3.897e-11) and library complexity (adjusted R-squared = 0.8938, p-value = 2.182e-14) before and after enrichment. The tests showed a strong positive correlation between the human DNA content of the pre-and post-enrichments, suggesting that an initial high human DNA content is a possible indicator of good target enrichment efficiency. In other words, libraries with low endogenous human DNA content may not enrich efficiently, even though the input DNA amount before the enrichment was high. Moreover, over-amplifying dsDNA libraries with low endogenous human DNA content led to higher PCR duplication rates.

We evaluated the enrichment efficiency of the Twist and Tweap protocols based on the human proportion and library complexity. We found that the enrichment efficiency using the Tweap protocol (adjusted R-squared = 0.06455, p-value = 0.1935) was not significantly associated with the endogenous human DNA content before the enrichment. In contrast, the Twist protocol showed a slightly stronger, albeit still weak, association (adjusted R-squared = 0.1844, p-value = 0.07049), suggesting that the initial DNA content may have a more significant influence on the enrichment efficiency. These findings imply that the Tweap protocol could be more robust for dsDNA libraries with lower library complexity.

### Validation of the target enrichment efficiency based on sequencing information

After processing the sequencing data, we investigated the proportion of human reads with high mapping quality (MapQ > 30), library complexity, PCR duplicates, and short read rates (< 35 bp) of the unique dsDNA libraries by comparing shotgun and enrichment sequencing data (Figure 2, Supplementary Data SD1).

**Figure 2.**
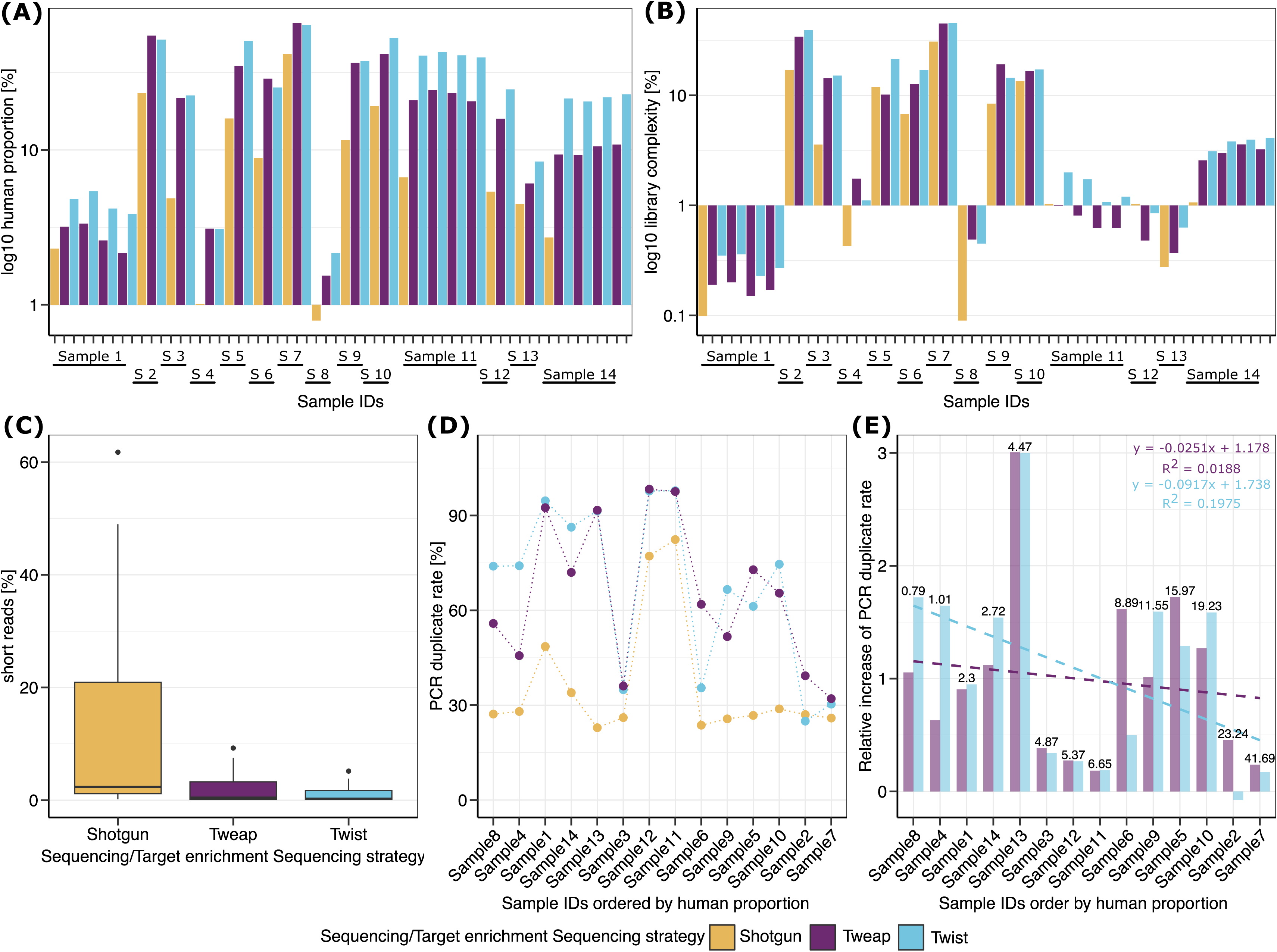
Overview of sequencing information obtained from the 14 dsDNA libraries selected for this study. **(A)** Barplot presenting the log10-transformed human proportion [%] of the shotgun and enrichment sequencing data. **(B)** Barplot presenting the log10-transformed library complexity [%] of the shotgun and enrichment sequencing data. **(C)** Short read rate distribution before and after the enrichment. **(D)** Scatterplot presenting the PCR duplicate rates of each sample ordered by the endogenous human DNA content before the enrichment. **(E)** A relative increase in the rate of PCR duplicates from each dsDNA library was achieved using the Tweap or Twist protocol ordered by the endogenous human DNA content before the enrichment (values above each bar). The dashed lines present the linear regression model fit to the relative increase rates. S = Sample ID.

The human proportion of the 14 selected dsDNA libraries were between 0.79 % and 41.69 % (mean = 10.63 %, median = 6.01 %) for the shotgun data and between 1.54 % and 66.07 % (mean [total] = 26.99 %, median [total] = 24.97 %, median [Tweap] = 25.30 %, median [Twist] = 24.97 %) for the enrichment data, indicating that the target enrichment of human DNA was efficient (Figure 2A, Supplementary Data SD1). The library complexity after enrichment increased from 6.84 % (median = 2.32 %) to 11.89 % (median = 6.06 %) on average (Figure 2A). In contrast, the PCR duplicate and short-read rates changed significantly after the enrichment. Here, the PCR duplicate rates for shotgun data were moderate (median = 27.03 %); however, after the enrichment, the majority of the dsDNA libraries showed a PCR duplicate increase of between 1 – 3 X, implying that especially enriched dsDNA libraries with three times more PCR duplicate rates were over-amplified (Figure 2C and 2D, Supplementary Table S5). Both protocols showed similar PCR duplicate rates (median [Tweap] = 63.67 %, median [Twist] = 74.03 %; linear regression model [total rates]: adjusted R-squared = 0.2552, p-value = 0.001205). Tweap protocol presented a lower increase of PCR duplicate rates than the Twist protocol in samples with low endogenous DNA content; however, a Welch two-test rejected this hypothesis (t = 0.23879, df = 25.603, p-value = 0.8132), suggesting that both protocols had produced a similarly high number of PCR duplicates. On the other hand, the short read rates of the dsDNA libraries decreased from 14.54 % (shotgun sequencing) to 1.63 % on average (mean [Tweap] = 2.10 %, mean [Twist] = 1.16 %) (Figure 2B) and the final average read length slightly increased from 82 (shotgun sequencing) to 93 bp (mean [Tweap] = 91 bp, mean [Twist] = 96 bp) after the enrichment.

### Estimation of the capture sensitivity and enrichment factor

We called the SNPs of the ‘1350k’ panel using the software ANGSD (Korneliussen, Albrechtsen and Nielsen, 2014a) and PLINK v1.9 (Chang *et al*., 2015a) and extracted SNPs included in the ‘1240k’ and ‘human origin’ (HO) panel (See Supplementary Material 1 for details, Supplementary Data SD1E). To investigate the efficiencies of the protocols, we calculated the capture sensitivity (CS) and enrichment factor (EF) of each sample using two methods: i) considering only the raw count of SNPs, and ii) normalising by the targeted reads and target/genome size, 102,263,103 and 3,053,308,597 (*f*_target space_ = 3.88×10^−4^), respectively (Hernandez-Rodriguez *et al*., 2018).

First, we focused on the enrichment factor EF_SNP-based_ (number of SNPs_post-capture_/number of SNPs_pre-capture_ (multi-pool samples were merged before calculation) and found that the EFs of the Twist protocol (median = 20.665) were higher than the Tweap protocol (median = 17.4) (See Table 1, Supplemental Figure S4, Supplementary Data SD1E), though a Pearson correlation test showed a statistically significant positive correlation (cor = 0.9766457, p-value = 2.228e-09) between the protocols (Bern, Walton-Day and Naftz, 2019). In addition, we investigated the correlation between the human proportion and library complexity, as well as the EF (Figure 3A and 3B). We found that an increase in the enrichment factor is not directly correlated with the human content of a selected dsDNA library (Figure 3C), opposing the hypothesis of a direct relation between a higher human content and successful SNP recovery (Supplementary Table S5).

**Figure 3.**
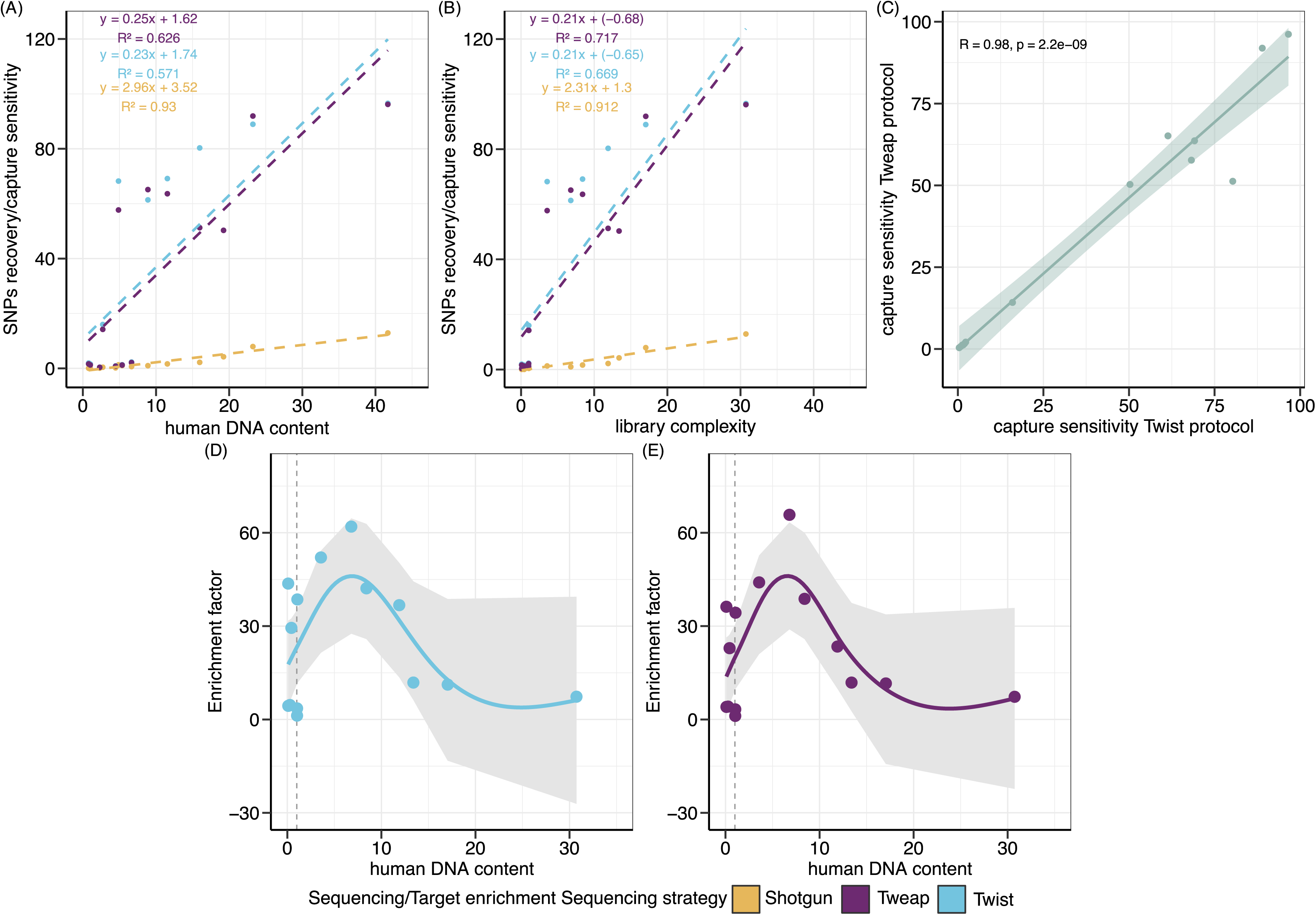
Visualisation of the correlation between the human proportion/library complexity and SNPs recovery/capture sensitivity for the shotgun data (yellow), Tweap (purple) and Twist (blue) protocols, and generalised additive modelling (GAM) fitted to the enrichment factor distribution. **(A/B)** Regression model prediction for all samples. **(C/D)** Regression model prediction excluding the samples below 1% endogenous human DNA content. The lines show the smooth effects of the endogenous human DNA content and the enrichment factor, with 95% confidence intervals presented in grey. The vertical dashed line presents the endogenous human DNA content threshold of 1%. (A) Scatterplot with regression lines to correlate the human proportion with the SNPs recovery/capture sensitivity. (B) Scatterplot with regression lines to correlate the library complexity with the SNPs recovery/capture sensitivity. (C) Correlation graph between the capture sensitivity between the Tweap and Twist protocol. (D/E) Regression model prediction of Twist and Tweap, respectively. The lines show the smooth effects of the human DNA content and the enrichment factor, with 95% confidence intervals presented in grey. The vertical dashed line presents the endogenous human DNA content threshold of 1%.

We further investigated this effect by applying a fitting generalised additive model (GAM) with a 95% confidence interval on the EF_SNP-based_ (Figure 3D and 3E, Supplementary Data SD1E). Here, both protocols have the highest enrichment factors for individuals with a human proportion between 1 % and 10 %. Interestingly, the enrichment factors dropped significantly when the human DNA content was above 10 %, suggesting that a range of 1 – 10 % has a significant potential to retrieve adequate SNPs.

Under consideration of the number of target reads and genome size, the Twist protocol (median = 0.264) presents overall better enrichment factors than the Tweap protocol (median = 0.15775), although both protocols present similar capture sensitivities (median [Tweap] = 32.245, median [Twist] = 33.135) (Supplementary Data SD1F). Thus, we applied a paired Wilcoxon signed-rank test to compare the enrichment factors and capture sensitivity (EF/CS of Tweap vs. Twist) to assess potential differences between the protocols. While the test for the EF indicated statistically significant differences (p-value = 0.03028), the test of the capture sensitivity showed only a modest but significant difference (p-value = 0.04944). While these tests indicated that both protocols are generally effective, the SNP recovery and efficiency of a target enrichment can greatly vary between the experiments and other factors, such as technical variabilities in the (sequencing) workflows.

To explore the possibility of enriching dsDNA libraries with ultra-low library complexity (<1%), we utilised multi-pool hybridisation reactions of samples 1, 11, and 14 (Supplementary Data SD1). Using Venn diagrams, we estimated the number of overlapping and non-overlapping ‘1350k’ SNPs to show the efficiency of multi-pooling (Figure 4). We found that both protocols presented different numbers of non-overlapping SNPs between the enriched dsDNA libraries, confirming the theory that the number of SNPs varies between dsDNA libraries. Notably, the enriched samples using the Tweap protocol (sample 1 = 90.46 %, sample 11 = 90.74 %, sample 14 = 81.93 %) showed a higher proportion of non-overlapping targeted SNPs between the different dsDNA libraries than the Twist protocol (sample 1 = 83 %, sample 11 = 82.22 %, sample 14 =76.84 %) suggesting that Tweap may enrich higher number of non-overlapping SNPs, which are essential for genome-wide SNP analysis.

**Figure 4.**
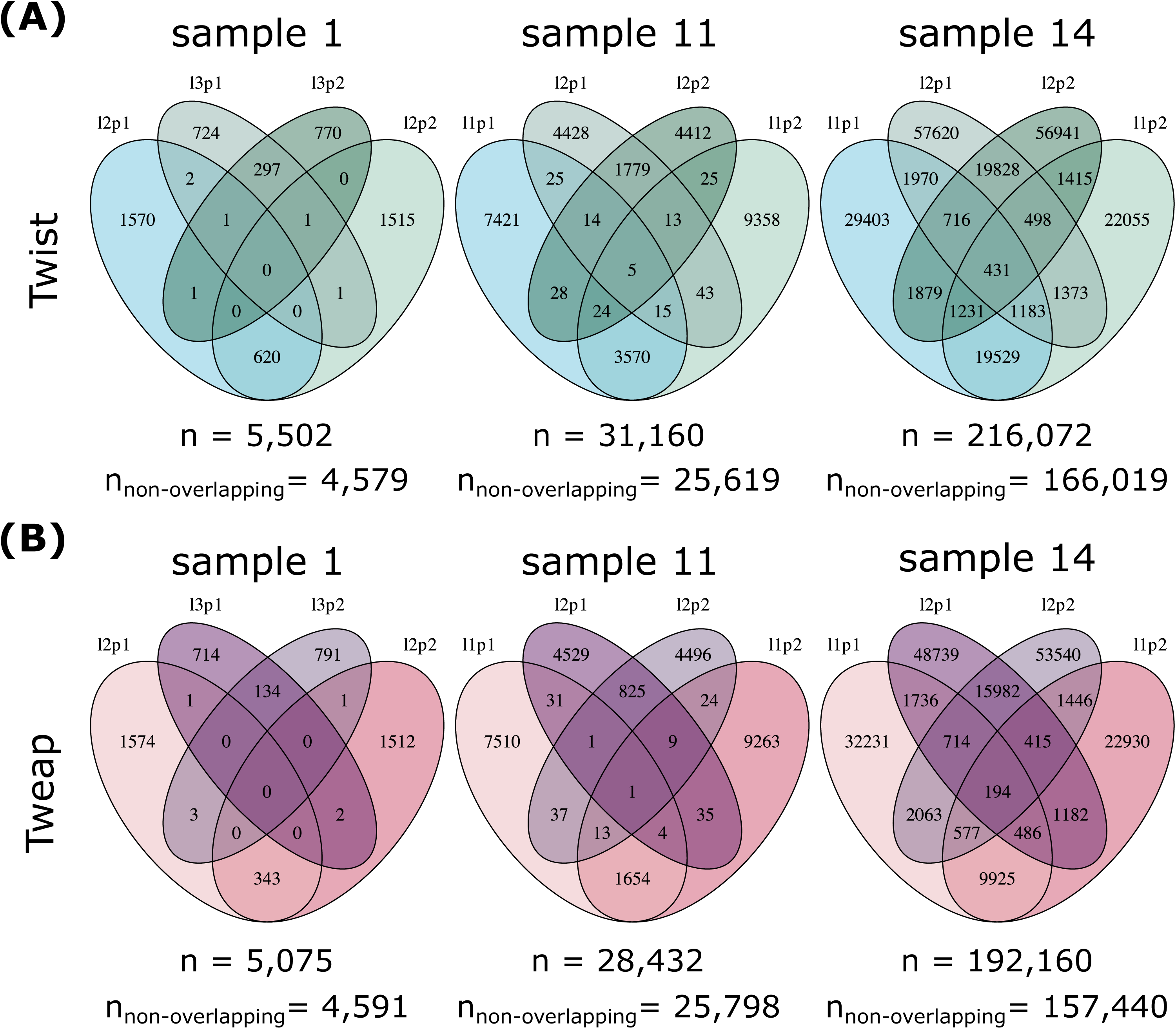
Venn diagrams displaying the number of overlapping and non-overlapping ‘1350k’ SNPs using multi-pooled dsDNA libraries for the target enrichment of samples 1, 11, and 14 with (A) the Twist protocol, and (B) the Tweap protocol. n = total number of SNPs. l = library. p = PCR. The numbers represent the number of SNPs before merging the single BAM files.

In addition, we also selected four samples (samples 1, 4, 8, and 13) with ultra-low library complexity (below 1 %), presenting an average of 0.224 % to investigate if the two protocols could enrich libraries with low human proportion as well as library complexity (Carpenter *et al*., 2013). Here, 2 out of the 4 selected dsDNA libraries (samples 4 and 8) had an outstanding enrichment factor of more than 20 X and 35 X and recovered more than 16,000 and 22,000 SNPs when using both protocols, respectively. This suggests the possibility of retrieving an adequate number of SNPs for genome-wide SNP analysis of individuals whose libraries contain low DNA yields (Supplementary Data SD1).

## Discussion

We have refined the employed protocols for target enrichment via in-solution hybridisation of DNA libraries and have successfully replaced the streptavidin-coated superparamagnetic beads required for the enrichment with the Dynabeads from Invitrogen. Following the replacement, we were able to enrich the dsDNA libraries using the Twist ‘Ancient Human DNA’ panel at a reduced cost. Throughout the study, both Twist and Tweap protocols were found to be effective in retrieving a sufficient number of SNPs required for downstream analysis of ancient individuals. The Twist protocol may seem to have a better performance in terms of presenting higher DNA concentrations after the enrichment and coverage depth of the target regions, leading to the retrieval of more SNPs; however, the duplicate rates were higher than Tweap. The Tweap protocol, on the other hand, was more efficient in capturing non-overlapping SNPs when performing a multi-pool hybridisation reaction (Figure 4).

The Twist protocol has been tested on single-stranded USER-treated and double-stranded partial-UDG-treated DNA libraries with endogenous human DNA content between 0.10 and ∼96 % (median = 68.66 %) (Rohland *et al*., 2022a).

Here, we performed the Twist and Tweap protocols on 14 blunt-end repaired dsDNA libraries of ancient individuals with endogenous human DNA content between 0.79 and 41.69 %. We decided to omit any additional treatment of the dsDNA libraries to ensure the presence of ancient DNA fragments, which are represented by the average deamination rates (mean [C>T] = 26.47 %, mean [G>A] = 26.37 %), and the average short read length (mean = 82 bp) before the target enrichment. After the enrichment, we see that the average read length distributions increased more for Twist than for Tweap, implying that the read length distribution is affected by the applied protocol. Others have reported similar findings and concluded that the shift in the average read length led to a higher retrieval of SNPs for downstream analysis (Rohland *et al*., 2022a).

Both protocols enriched effectively up to 60X when the human proportion was between 1 % and 10 %. However, for libraries with more than 10 % human proportion, the enrichment factors showed a steep decline. We also explored the presumption that enrichments of ancient DNA libraries (n = 4) with low human content would not hold sufficient SNPs for broad-scale population genetic analysis (Cruz-Dávalos *et al*., 2017, 2018; Yaka *et al*., 2024) (Figures 1 and 4). Interestingly, two of the four samples showed sufficient enrichment, indicating that the selection of target enrichment is not only influenced by the human proportion, library complexity, PCR duplicate, or short read rates, but also subject to inherent randomness. However, exploring possible factors further was beyond the scope of this study due to the limited number of dsDNA libraries available.

Several target enrichment studies have previously reported increasing PCR duplicate rates relative to shotgun screening data (Rohland *et al*., 2022a; Yaka *et al*., 2024). Our study showed similar findings when applying both protocols. However, the Tweap protocol had lower PCR duplicate rates than the Twist protocol when applied to dsDNA libraries, especially when the human proportion was between <1 % and 3 % before the enrichment. Concerning aDNA studies, a low number of PCR duplicates is preferred to increase the human coverage of the individuals by performing additional sequencing runs of the same dsDNA libraries. The high amount of PCR duplicates would interfere with more unique DNA sequencing reads required for downstream analysis. Throughout the optimisation of both protocols, we frequently observed the presence of ‘bubble DNA’ artefacts, later associated with the high PCR duplicates after sequencing. Therefore, we successfully applied a reconditioning PCR amplification method on most of the enriched dsDNA libraries to reduce the ‘bubble DNA’ before sequencing.

This step can help to minimise overamplified or mispaired molecules by using a one-cycle PCR amplification without a hold to cool down (Thompson, Marcelino and Polz, 2002a). Interestingly, dsDNA libraries enriched using the Tweap protocol displayed minimal to no visible ‘bubble DNA’ artefacts, suggesting a lower occurrence of PCR-induced structural anomalies, leading to a higher input volume for sequencing. (Supplemental Figure S3).

One of our aims was to reduce the costly reagents required for the target enrichment of ancient DNA libraries by replacing the beads for capturing the biotinylated probes bound to the targeted DNA and purifying the amplified dsDNA fragments. By using the Dynabeads for Target Enrichment, we reduced the purchase of additional Twist kits, allowing researchers to reallocate their funding (Table 1). For an estimation of the costs, we calculated the theoretical number of enrichments using the 2, 12, and 96-reaction kits for the Twist ‘ancient human DNA’ panel (input: 1 µl) and ‘mtDNA’ panel (input: 0.167 µl), resulting in 8, 48, and 384 reactions for the ‘ancient human DNA’ panel and 48, 287, and 2,299 reactions for the ‘mtDNA’ panel. Then, we estimated the price for one reaction using the price lists specified for Northern European countries. Our final estimated costs (excluding the panels) of performing one enrichment reaction using the Twist protocol and Tweap protocol were $27 USD and $5 USD, respectively. Concluding a potential savings of more than $20 USD per hybridisation reaction when using the Tweap protocol.

Finally, the environmental impact should be considered when working in a life science laboratory. For a user-friendly performance of the target enrichment, Twist Bioscience provides the binding and washing buffers in a kit; however, for the modified Twist protocol applicable for dsDNA with low endogenous human DNA content, more binding buffer is required to bind the streptavidin binding beads, leaving a high amount of washing buffers behind. Researchers would discard those reagents and single-use plastic bottles at the end of the experiment, contributing to both chemical and plastic waste. In contrast, the Tweap protocol only requires reagents for the preparation of the binding & washing buffer, which are commonly used in well-equipped life science laboratories and hence, build a foundation for sustainability in scientific research.

## Conclusion

In conclusion, our Tweap protocol presents a potential alternative for the target enrichment of aDNA using the Twist ‘Ancient Human DNA’ panel, allowing researchers to increase the number of ancient individuals for their studies, without compromising capture efficiency, by reducing costs for the reagents and supporting more sustainable research practices. Based on this study and additional enriched dsDNA libraries using the Tweap protocol in our aDNA facility at Uppsala University (Sweden), we showed that multi-pool hybridisation reactions of poorly preserved samples can still yield sufficient SNPs for broad-scale population genetic analysis. We believe that this approach could enable research in previously inaccessible geographical regions and allow the inclusion of individuals with limited human content who would otherwise have been excluded from population-scale studies.

## Declarations

### Ethical statement

The ancient human skeletal remains included in this study are a continuation of the ERC Advanced Grant: ‘The Enigma of the Hyksos’ (668640) led by Manfred Bietak and Holger Schutkowski from 2016 to 2021. Collections curators and archaeologists have been involved throughout the projects and continue to be collaborators and advisors. We sampled individuals with multiple elements present, ideally an antimeres to the sample taken (when possible) and took detailed photographs of the material before performing the micro-destructive sampling. Furthermore, we sampled the minimum amount necessary for DNA recovery and used protocols that allow for different types of biomolecular or chemical analysis (simultaneously or in the future) to be carried out on the same bone material. The DNA extractions are currently stored either at the aDNA facility at Uppsala University (Sweden) or the University of Tartu (Estonia). The prepared and sequenced dsDNA libraries are stored at Uppsala University.

### Availability of data and materials

The datasets used and/or analysed during the current study are available from the corresponding author on reasonable request.

### Authors’ contributions

C.B., M.J., and T.S. designed the study. C.L.S. and H.N. extracted the DNA of the human skeletal remains at the Institute of Genomics, University of Tartu (Estonia). M.O.G. and T.S. performed the laboratory work at Uppsala University (Sweden), and E.T. provided technical assistance. C.B. performed the raw data processing of the shotgun sequencing data. C.B. and T.S. conducted the statistical analysis of the data. T.S. wrote the first draft of the manuscript with input from C.B. and E.T. All authors reviewed and accepted the manuscript.

## Acknowledgements

We thank the archaeologists for the permission to extract DNA from human skeletal material used in this study and to test our refined protocol on those samples. Sequencing was performed by the SNP&SEQ Technology Platform in Uppsala. The facility is part of the National Genomics Infrastructure (NGI) Sweden and the Science for Life Laboratory. The Swedish Research Council and the Knut and Alice Wallenberg Foundation also support the SNP&SEQ Platform. Ancient DNA data analysis was supported by the SciLifeLab Ancient DNA unit, funded by the Science for Life Laboratory and the Swedish Research Council. The data handling and computations were enabled by resources provided by the National Academic Infrastructure for Supercomputing in Sweden (NAISS) at Uppmax, partially funded by the Swedish Research Council through grant agreement no. 2022-06725. We thank Thijessen Naidoo for processing the enrichment sequencing data. T.S. thanks Nadin Rohland for advice with the Twist Ancient Human Panel protocol. We thank Andrea Soler i Núñez for preparing 2 library samples for her MA thesis. T.S. thanks BTS for their music and the colour palette used in this publication. We thank E. Israel Chávez Barreto for his illustration of Supplementary Figure S2. The authors acknowledge using OpenAI’s ChatGPT for assistance in refining the language.

## Supplementary Material 1

### Materials and Methods

#### Ancient DNA Laboratory Workflow

##### Sampling and DNA Extraction

The sampling and DNA extraction of human skeletal remains of ancient individuals (petrous bone = 10, tooth = 4, ∑ = 14) were conducted by Christiana L. Scheib and Helja Kabral at the Institute of Genomics, University of Tartu (Estonia) (Keller, 2021b, 2021a, 2022, 2023) except sample 1 (Table 1, Supplementary Data SD1). Sample 1 was sampled at the Institute of Genomics, University of Tartu (Estonia), and the extraction was performed in the aDNA unit of the Human Evolution program following a pre-digesting powder protocol with an incubation time of 30 min (Damgaard *et al*., 2015). One extraction was performed per sample for this project and eluted in 100 µl pre-heated elution buffer (EB buffer) at 37 °C provided in the MinElute PCR Purification Kit (QIAGEN #28006). The DNA concentrations of the extracts were measured using a Qubit 2.0 Fluorometer to evaluate the presence of DNA molecules. The DNA concentrations varied between <0.05 (too low to be measured) and 4 ng/µl. In 2023, the remaining part of the extracts (20 - 100 µl) were sent to the Human Evolution program, Department of Organismal Biology, Uppsala University (Sweden), for further analyses using target enrichment via in-solution hybridisation.

Before the library preparation, the DNA extracts were cleaned using the OneStep PCR Inhibitor Removal Kit (Zymo Research #D6030) to remove any contaminants that could inhibit DNA sequencing preparations efficiently. We added 600 µl of the preparation solution to each Zymo-Spin™ III-HRC column and centrifuged it at 8,000 rcf for 3 min. After placing the column into a new clean 1.5 ml Eppendorf® LoBind microcentrifuge tube, 20 – 100 µl of DNA extract was added and centrifuged at 16,000 rcf for 3 min. The column was removed from the 1.5 ml Eppendorf® LoBind microcentrifuge tube and the ‘cleaned’ DNA extracts were stored at -20 °C.

##### Library Preparation

Blunt-end repaired, double-stranded DNA (dsDNA) and double-indexed libraries, with unique P5 and P7 index combinations, were built at the Ancient DNA facility, Uppsala University (Sweden). The dsDNA libraries and library blanks (starting input per reaction: 20 µl DNA or nuclease-free molecular biology-grade water) were built following a modified version of the established protocol ‘Illumina Sequencing Library Preparation for Highly Multiplexed Target Capture and Sequencing’ (Meyer and Kircher, 2010a), where the MinElute PCR Purification Kit was used, instead of SPRI beads, to clean up the reactions, following manufacturer’s instructions. After the adapter fill-in step, the libraries were stored at -20 °C.

##### Estimation of amplification cycles before dual-indexing

We used a quantitative PCR (qPCR) method to estimate the number of cycles needed before dual indexing to optimise the number of unique indexed fragments while minimising the number of PCR duplicates in dsDNA libraries. For the qPCR, we used 12.5 µl of the ready-to-use solution Maxima SYBR Green/ROX qPCR Master Mix (Thermo Scientific #K0221), 10.5 µl of *dd*H_2_O, and 0.5 µl IS7_short_amp.P5 (ACA CTC TTT CCC TAC ACG AC) and 0.5 µl IS8_short_amp.P7 (GTG ACT GGA GTT CAG ACG TGT) primers, and 1 µl DNA library per reaction, in duplicates. The qPCR was performed on a BioRad CFX96 using the qPCR program presented in Supplementary Table S1. The C_t_ values of the dsDNA libraries were used to determine the number of cycles for dual indexing of each dsDNA library by taking the mean of the duplicates and adding 2 cycles to them (Supplementary Data SD1).

**Supplementary Table S1.** qPCR program to estimate the number of cycles.

|  | Temperature (°C) | Time | Cycle(s) |
| --- | --- | --- | --- |
| Incubation | 95 | 10 min |  |
| Denaturation | 95 | 30 sec | 32 |
| Annealing | 60 | 30 sec |  |
| Extension | 72 | 30 sec |  |

##### Double-indexing of the dsDNA libraries using amplification

To prepare the dsDNA libraries for sequencing, we used unique double-index combinations added during the amplification of the dsDNA libraries and library blanks in duplicates. We included one additional PCR blank with a unique double-index combination per cycle. For the amplification, we prepared a master mix with 5 µl of Taq Gold buffer (10 x), 5 µl of MgCl_2_, 0.5 µl of dNTPs (25 mM each), 1 µl of AmpliTaq Gold DNA Polymerase, and 30.5 µl of molecular biology grade nuclease-free water. We added 6 µl of dsDNA library or molecular biology grade nuclease-free water per reaction and 2 µl of pre-combined P5 and P7 index primers (each 10 mM), reaching an amount of 50 µl. Each PCR reaction was set up with the number of cycles estimated from the qPCR (see the above section), but with a maximum of 20 cycles (Supplementary Table S2).

**Supplementary Table S2.** Amplification program for blunt-end repaired dsDNA libraries.

|  | Temperature (°C) | Time | Cycle(s) |
| --- | --- | --- | --- |
| Initialisation | 94 | 12 min |  |
| Denaturation | 94 | 30 sec | number of cycles<br>$= \bar{x} C_t + 2$ |
| Annealing | 60 | 30 sec |  |
| Extension | 72 | 45 sec |  |
| Final extension | 72 | 10 min |  |
| Hold | 12 | ∞ |  |

The PCR products were cleaned using the MinElute PCR Purification Kit or AMPure XP Bead-based reagents (Beckman Coulter #A63881) following a modified protocol for short DNA fragments as present in ancient DNA libraries. With the MinElute PCR Purification Kit, 500 µl of PB buffer and the first duplicate were added to a MinElute column. The MinElute column was spun down at 13,000 rpm for 1 min. The second duplicate was added to the column with 500 µl of PB buffer. The MinElute column was spun down again and placed into a new collection tube, and then 650 µl of PE buffer was added to the column. After the spin down, the MinElute column was put into a new collection tube, and a final dry-down spin of 1 min was performed. The MinElute column was placed in a new 1.5 ml Eppendorf® LoBind microcentrifuge tube, and the libraries were eluted in 35 µl of EB buffer. The tubes were incubated at room temperature for 10 min. After a spin-down at 13,000 rpm for 2 min, the MinElute columns were discarded, and the double-indexed dsDNA libraries were stored at -20 °C.

Using the AMPure XP Bead-based reagent, the duplicates of each dsDNA library were pooled together into a new 1.5 ml Eppendorf® LoBind microcentrifuge tube, and 1.8x volume of the AMPure XP beads was added. The tubes containing the dsDNA libraries were vortexed, pulse spun down and incubated for 5 min at room temperature. Then, the tubes with a slightly open lid were placed on a magnetic rack for 5 min or until the solution was clear at room temperature. The 1.5 ml Eppendorf® LoBind microcentrifuge tubes stayed on the magnetic rack throughout the purification procedure. After the incubation, the supernatant was removed without disturbing the bead pellet. Then, 250 µl of 70 % ethanol was added to the tubes, incubated for 1 min on the magnetic rack, and the ethanol was discarded. This washing step was repeated two more times. Then the bead pellets were air-dried with an open lid on the magnetic rack until all ethanol was evaporated (around 5 – 10 min). The tubes were removed from the magnetic rack and 40 µl of EB buffer was added to each tube while washing the beads down from the wall. After the incubation of 10 min, the tubes were placed back on the magnetic rack, and 37 µl of EB buffer was taken from the supernatant, which contained the eluted DNA fragments. The final dsDNA libraries were stored at -20 °C.

#### Target enrichment via in-solution hybridisation

To develop and test the efficiency of the Tweap protocol, we performed the Twist ‘Twist Target Enrichment Standard Hybridization v1’ protocol (https://www.twistbioscience.com/resources/protocol/twist-target-enrichment-standard-hybridization-v1-protocol, Revision 4.0) and Tweap protocols (Figure 1) on a small set of 14 samples (11 single-pool and three multi-pool samples (Supplementary Figure S3)), applying the same starting conditions, such as DNA starting concentration, input volume, and the volume of the streptavidin binding beads (Supplementary Data SD1).

##### Preparation of the selected libraries

Twist Bioscience recommends a starting concentration between 500–1000 ng per reaction for successful target enrichment of ancient DNA libraries. We used the Equinox Library Amplification Kit w/o amplification primers (Watchmaker Genomics #7K0021-096) to re-amplify selected dsDNA libraries of low DNA concentrations to maximise the presence of the unique target DNA fragments (Supplementary Data SD1). We estimated the number of PCR cycles based on the number of DNA copies calculated using the DNA concentration (measured with Qubit 3) and the average DNA fragment length (analysed with the 4200 TapeStation instrument) (Supplementary Data SD1). We prepared a 40/45 µl amplification reaction mix per reaction (input: 10/5 µl respectively) containing 12.50/17.50 µl nuclease-free molecular biology-grade water, 25 µl Equinox Library Amplification Mix (final concentration: 1 x), and 2.5 µl of customised Amplification primers acquired from Integrated DNA Technologies (IDT) (sequencing of IS5_reamp.P5: AAT GAT ACG GCG ACC ACC GA and IS6_reamp.P7: CAA GCA GAA GAC GGC ATA CGA, final concentration: 10 µM) (Meyer and Kircher, 2010a), and amplified the libraries using 5 or 6 PCR cycles to maximise coverage uniformity and fidelity (Supplementary Table 3).

**Supplementary Table S3.** Amplification program for the re-amplification of dsDNA libraries.

|  | Temperature (°C) | Time | Cycle(s) |
| --- | --- | --- | --- |
| Initialisation | 98 | 45 sec |  |
| Denaturation | 98 | 15 sec | 5 or 6 |
| Annealing | 60 | 30 sec |  |
| Extension | 72 | 30 sec |  |
| Final extension | 72 | 1 min |  |
| Hold | 12 | ∞ | |

The re-amplified dsDNA libraries were cleaned with the DNA Purification Beads included in the Twist Dry Down Beads kits (Twist Bioscience, part number #104325) following the clean-up steps provided in the ‘Twist Target Enrichment Standard Hybridization v1’ protocol. After the Beads-drying step on the magnetic rack at room temperature, we eluted the DNA in 40 µl EB Buffer and transferred 35 µl into a new clean 1.5 ml Eppendorf® LoBind tube.

After the amplification, we pooled the re-amplification batches of one dsDNA library and measured the final concentration (Supplementary Data SD1). The re-amplified dsDNA libraries were stored at -20 °C. To minimise the PCR duplicate rate, we increased the starting volume to reach the minimum target concentration of 500 ng, except for samples 3, 7, and 13, where the DNA concentrations of the original dsDNA libraries were low. To incubate the dsDNA libraries for the hybridisation reaction, we eluted the dsDNA libraries in 10 µl Blocking Solution Master Mix following the ‘Alternate pre-hybridisation DNA concentration protocol’ appended to the Twist Bioscience protocol.

##### Twist Target Enrichment Protocol

The Twist protocol was performed according to Rohland *et al*. 2022, which was based on the ‘Twist Target Enrichment Standard Hybridization v1’ protocol with modifications (Rohland *et al*., 2022b). The enriched dsDNA libraries were stored at -20 °C (Supplementary Data SD1).

##### Tweap Target Enrichment Protocol

To perform the target enrichment for aDNA libraries, we also based our alterations on the modified ‘Twist Target Enrichment Standard Hybridization v1’ protocol (Figure 1). We replaced the following reagents: i) Twist streptavidin binding beads with Dynabeads™ Streptavidin for Target Enrichment (Invitrogen #65606D); ii) Twist DNA Purification beads with AMPure XP Bead-Based Reagent (Beckman Coulter #A63881); and iii) ‘Twist Hybridisation & Wash Buffers’ kit with a home-made 2X Binding and Washing (B&W) buffer (10 mM Tris-HCl pH 7.4, 1 mM EDTA, 2 M NaCl). The enriched dsDNA libraries were stored at -20 °C. Detailed steps of the ‘Tweap Target Enrichment Hybridisation’ protocol are available on protocols.io (https://www.protocols.io/blind/2107D8C61E8E11F0A7670A58A9FEAC02), where we included a list of all required reagents, equipment and parameter settings.

##### Post-capture amplification

For the post-capture amplification, we used the standard amplification protocol described in the ‘Twist Target Enrichment Standard Hybridization v1’ protocol, using the Is5_reamp.P5 and Is6_reamp.P7 primers included in the Twist Hybridisation and Wash Kit or from IDT. We used 22.5 µl of the enriched dsDNA library for the amplification. We varied the number of cycles for the amplification between 10 and 23 cycles (Supplementary data SD1). The number of cycles was determined depending on the DNA starting concentration and the endogenous human DNA content of the shotgun library. The enriched dsDNA libraries were cleaned using the Twist DNA Purification Beads for the Twist protocol or AMPure XP beads for the Tweap protocol.

##### Quality and Quantity Assessment of the dsDNA libraries pre- and post-hybridisation

Two verification steps were implemented to validate the library preparation and measure the concentration of dsDNA libraries - i) fluorometric quantitation (Qubit® 3.0 Fluorometer, Invitrogen) to estimate the DNA concentration; and ii) parallel capillary electrophoresis (4200 TapeStation, Agilent) to evaluate the quality of the DNA libraries, using High Sensitivity D1000 ScreenTape (Agilent #5067-5584) or the D1000 ScreenTape (Agilent #5067-5582) according to manufacturers’ instructions (Supplementary Data SD1).

##### Troubleshooting

During the quality and quantity assessment of the enriched dsDNA libraries, multiple libraries presented high DNA concentrations, however, the qualitative analysis using a TapeStation Instrument showed a secondary peak right behind the 180 – 220 bp peak, which is usually the standard distribution of DNA fragments from aDNA libraries, or close to the upper marker at 1000 bp suggesting the presence of so-called ‘Bubble DNA’. Whereas the company Illumina excluded possible disruption during the sequencing process, as the dsDNA will be denatured, we decided to use a reconditioning PCR to decrease the size of the ‘Bubble DNA’ when the secondary peak was higher than that of the original dsDNA library. This application would break down the Bubble DNA using a modified version of a standard DNA amplification and minimise the possibility of off-target sequences during the hybridisation reaction. After the amplification, the libraries can be cleaned following the standard DNA purification protocol. In addition, to avoid generating ‘Bubble DNA’, we used fewer amplification cycles (10 or 15) for dsDNA libraries with initial high endogenous human DNA content (Supplementary figure S4).

##### Reconditioning DNA amplification

In addition to the lower number of PCR cycles, we applied a reconditioning DNA amplification on enriched dsDNA libraries with a large ‘Bubble DNA’ using a modified amplification protocol of the Equinox Library Amplification Kit w/o amplification primers (Supplementary figure S1) (Thompson, Marcelino and Polz, 2002b; Buchner, 2024). We used 5 – 10 µl of the enriched dsDNA library to enhance the DNA polymerase and ran a modified PCR program without hold (Supplementary Table 4). The reconditioned dsDNA libraries were stored at -20 °C and were ready to be pooled for sequencing.

**Supplementary Table S4.** Amplification program for the re-conditioning PCR.

|  | Temperature (°C) | Time | Cycle(s) |
| --- | --- | --- | --- |
| Initialisation | 98 | 45 sec |  |
| Denaturation | 98 | 15 sec | 1 |
| Annealing | 60 | 30 sec |  |
| Extension | 72 | 30 sec |  |
| Final extension | 72 | 1 min |  |

##### DNA sequencing

Before the enrichment, the double-indexed dsDNA libraries were shotgun-screened on an Illumina NovaSeq 6000 Sequencing system or a NovaSeq X Plus Sequencing system. The enriched dsDNA libraries were sequenced on a NovaSeq X Plus Sequencing system. For the direct correlation, the enriched dsDNA libraries were equal molar pooled, leading to the same pool concentrations and sequenced on the same sequencing platform. Sequencing was performed by the National Genomics Infrastructure, SNP&SEQ Technology Platform, Uppsala Biomedical Centre, Uppsala University (Sweden).

#### Raw Data Processing

##### Alignment to the reference genome

The raw sequencing data was processed at the Human Evolution Program, Uppsala University (Sweden), using an in-house computational pipeline developed for aDNA analysis. Before the alignment to the human reference genome (hs37d5), the residual adapter sequences and terminal consecutive G’s were removed using Cutadapt v. 2.3 (Martin, 2011), with the “--nextseq-trim 15” flag and the paired-end reads were merged using FLASH v 1.2.11 (Magoč and Salzberg, 2011) requiring a minimum overlap of 11 bases. The merged reads were aligned to the reference genome using Burrow-Wheeler Aligner (BWA) (Li and Durbin, 2009) *aln* v 0.7.17 with the parameters *-l 16500 -n 0.01 -o 2* and converted to a single-end alignment file using BWA *samse.* The files in sequencing alignment/map (SAM) format were then saved to binary alignment map (BAM) files using SAMtools v1.10 (Li *et al*., 2009). The PCR duplicates were removed by collapsing reads with identical start and end positions using a modified version of FilterUniqSAMCons_cc.py (Kircher, 2012). Finally, sequencing reads with less than 35 bp length and consensus below 90% to the human reference genomes were filtered out using a custom script. We used the same in-house computational pipeline for the shotgun and enriched dsDNA libraries (Supplementary Data SD1). For the statistical analysis, we excluded reads with low mapping quality (<30 bp) as they are excluded from the downstream analysis.

##### Sequencing statistics and aDNA authentication

To evaluate the ancient authenticity of the sequenced reads from the dsDNA libraries, we used SAMtools (Li *et al*., 2009) to estimate the aligned reads at various alignment stages (Supplementary Data SD1). The average read length was assessed by measuring the read lengths of randomly selected reads (maximum 2 million). The average read lengths for the shotgun-screened dsDNA libraries were between 50 - 110 bp (median = 81) and for the enriched dsDNA libraries between 50 - 130 bp (median = 89), showing overall an improvement of the average read length after the enrichment (Ginolhac *et al*., 2011b; Jónsson *et al*., 2013b). The deamination pattern of the shotgun dsDNA libraries was estimated by counting the presence of C→T substitutions at the 5’-end and G→A substitutions at the 3’-end for the first 25 bp (Supplementary Figure S2). The modern human DNA contamination rates were estimated on the mitochondrial DNA of the dsDNA libraries using the tool ContamMix (Fu *et al*., 2014). We estimated the depth and average coverage of the unique reads overlapping the Single Nucleotide Polymorphisms (SNPs) on the ‘1350k’ array (https://genome.cshlp.org/content/32/11-12/2068/suppl/DC1) (Rohland *et al*., 2022b) using the software Qualimap (García-Alcalde *et al*., 2012; Okonechnikov, Conesa and García-Alcalde, 2016).

##### Calling of Single Nucleotide Polymorphisms

Base quality recalibration was performed on the 10 outermost bases of each read by reducing the quality of all T’s on the 5’ end and all A’s on the 3’ end to Phred score 2 (#). The software ANGSD (Korneliussen, Albrechtsen and Nielsen, 2014b) was used to call the SNPs on the ‘1350k’ array (including autosomal and sex chromosomes) (*-doMajorMinor, doHaploCall*) applied on all variants (*-doCounts*). The output in FASTA format was converted to PLINK bed format via haploToPlink and Plink v1.9 (Chang *et al*., 2015b). The SNPs on the ‘1240k’ and the human origin (HO) arrays were extracted using Plink v1.9 (Table 1, Supplementary Data SD1).

##### Calculation of the enrichment factor

We calculated the enrichment factor using the number of biallelic SNPs on the ‘1350k’, ‘1240k’, and ‘HO’ arrays obtained from the shotgun and enrichment data as follows: 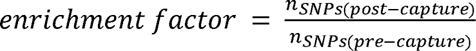 (Supplementary Data SD1).

##### Visualisation of the statistical analysis

The statistical analysis and visualisation of the data were performed using R 4.3.2 (R Core Team, 2023) and the following packages: *ggplot2, reshape2, tidyr, dplyr,* and *cowplot*. We plotted the *enrichment factor vs. the endogenous human DNA content of the shotgun data* and fitted a generalised additive model using the R package *mgcv* (https://cran.r-project.org/web/packages/mgcv/index.html) (Figure 2). The Venn diagrams were generated using the R package *VennDiagram* (Figure 3). To compare various parameters, we used a linear regression model function (lm()) provided by R.

## Supplementary Tables

**Supplementary Table S5.**
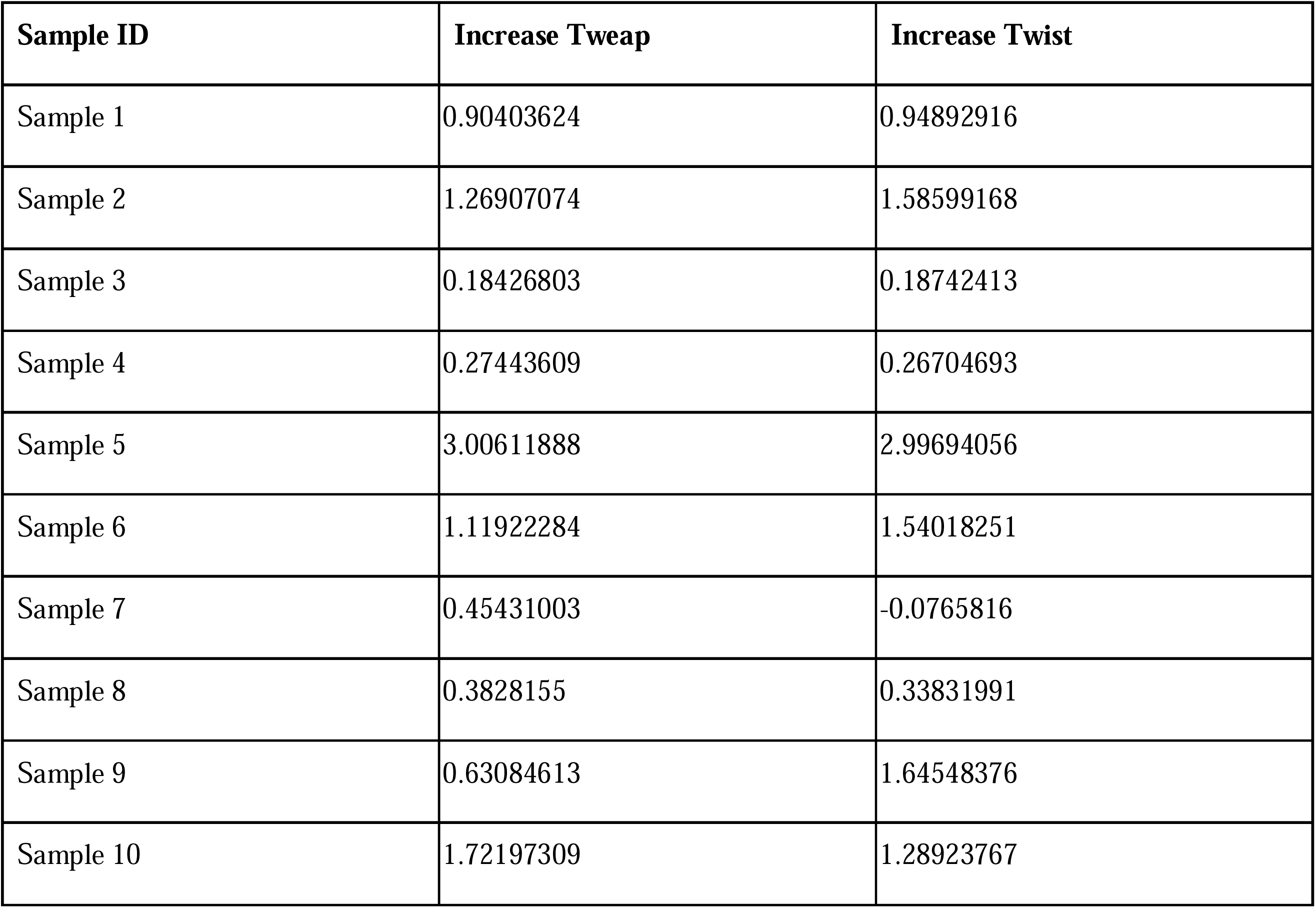
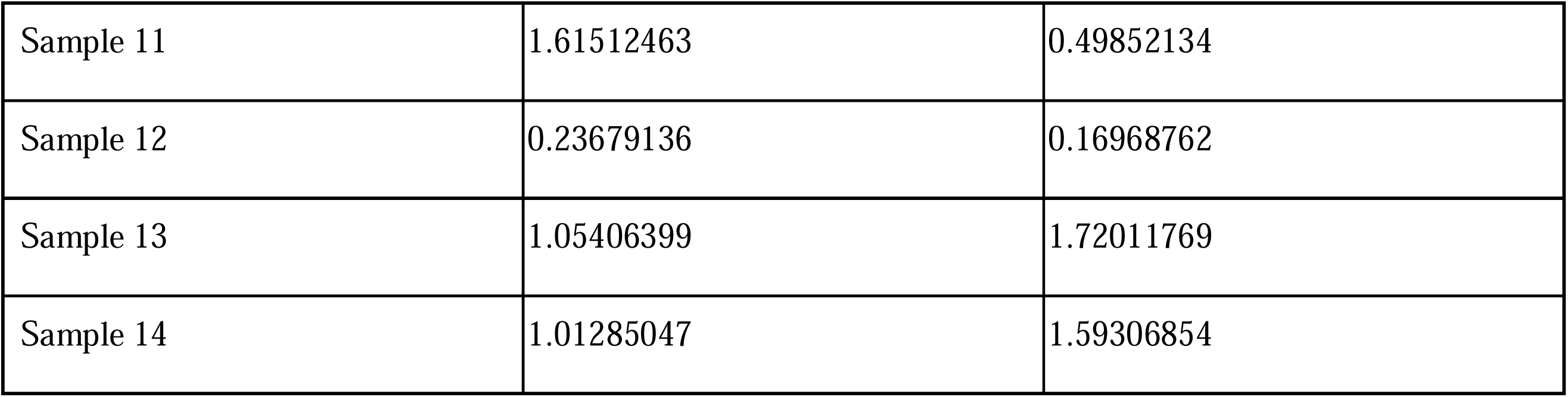
Calculated PCR duplicate increase rates after the enrichment.

| Sample ID | Increase Tweap | Increase Twist |
| --- | --- | --- |
| Sample 1 | 0.90403624 | 0.94892916 |
| Sample 2 | 1.26907074 | 1.58599168 |
| Sample 3 | 0.18426803 | 0.18742413 |
| Sample 4 | 0.27443609 | 0.26704693 |
| Sample 5 | 3.00611888 | 2.99694056 |
| Sample 6 | 1.11922284 | 1.54018251 |
| Sample 7 | 0.45431003 | -0.0765816 |
| Sample 8 | 0.3828155 | 0.33831991 |
| Sample 9 | 0.63084613 | 1.64548376 |
| Sample 10 | 1.72197309 | 1.28923767 |
| Sample 11 | 1.61512463 | 0.49852134 |
| Sample 12 | 0.23679136 | 0.16968762 |
| Sample 13 | 1.05406399 | 1.72011769 |
| Sample 14 | 1.01285047 | 1.59306854 |

**Supplementary Table S6.** Pearson’s correlation tests of capture sensitivity.

| Test | Correlation coefficient | p-value |
| --- | --- | --- |
| human proportion & capture sensitivity [Tweap] | 0.8089388 | 0.0004589 |
| human proportion & capture sensitivity [Twist] | 0.7772364 | 0.001071 |
| library complexity ~ capture sensitivity [Tweap] | 0.833533 | 0.0002125 |
| library complexity ~ capture sensitivity [Twist] | 0.859532 | 8.138e-05 |

## Supplementary Figures

**Supplementary Figure S1.**
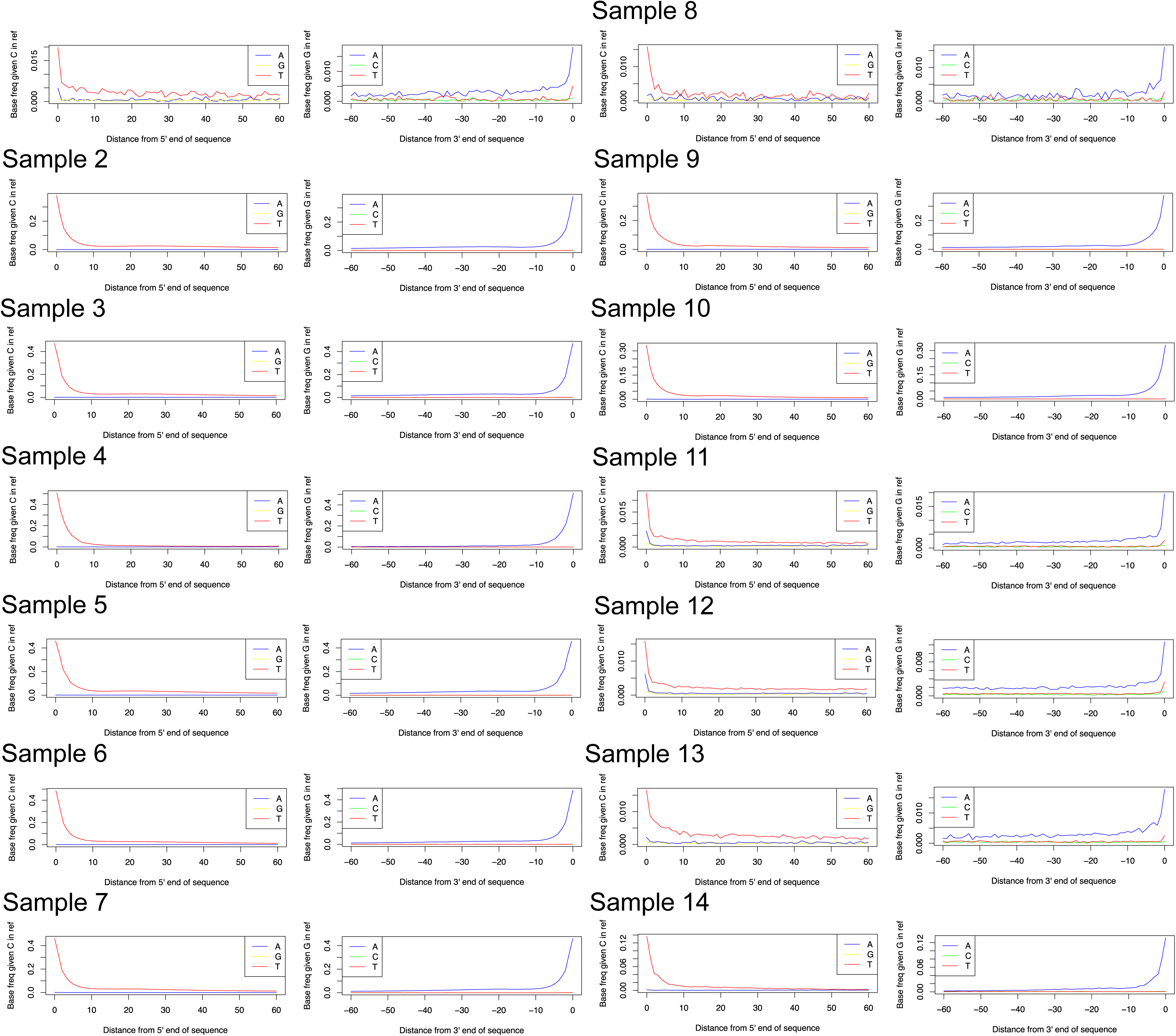
Deamination profiles of the selected dsDNA libraries after shotgun sequencing.

**Supplementary Figure S2.**
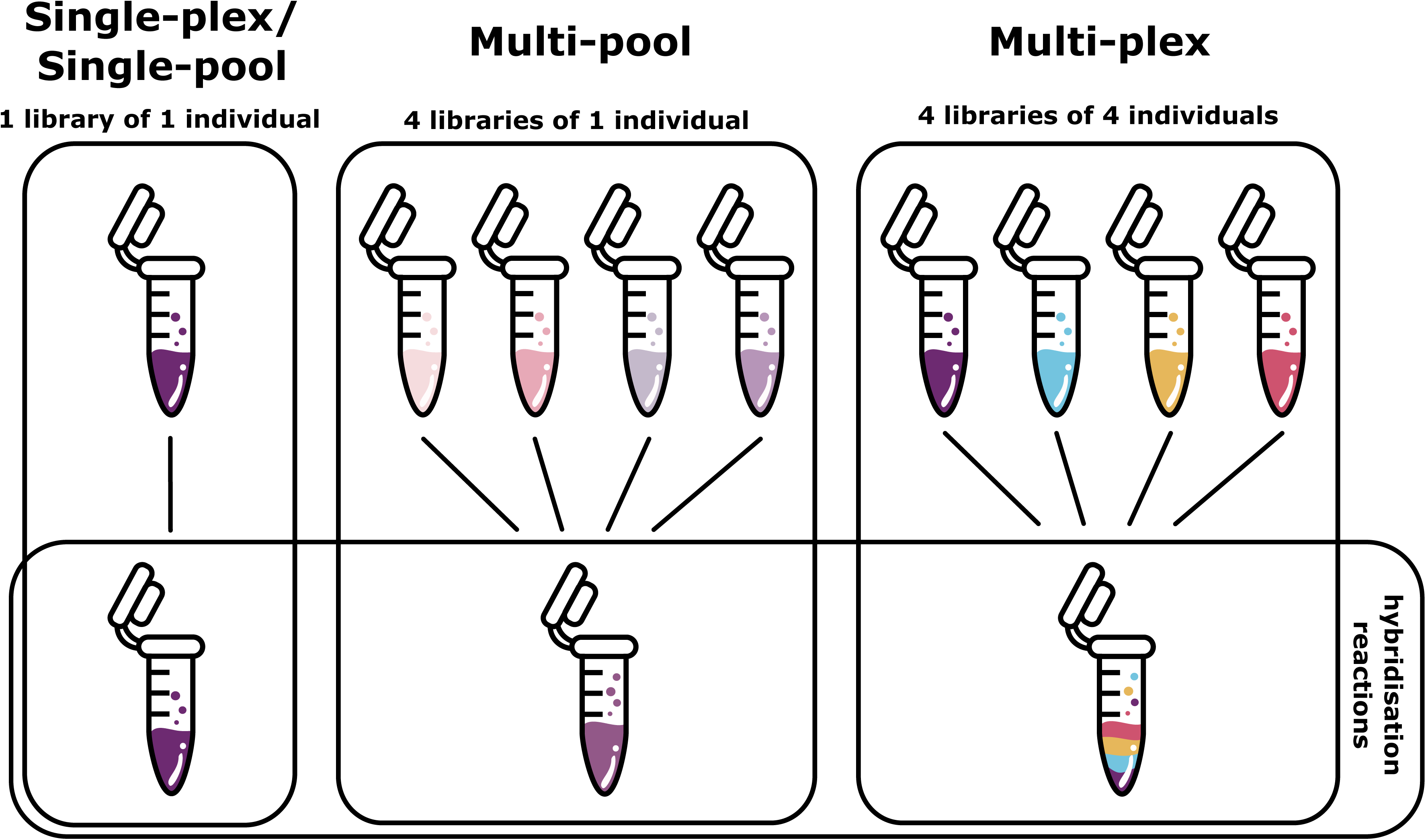
Graphical explanation of single- and multi-pool hybridisation reactions used in this study. Illustration by E. Israel Chávez Barreto.

**Supplementary Figure S3.**
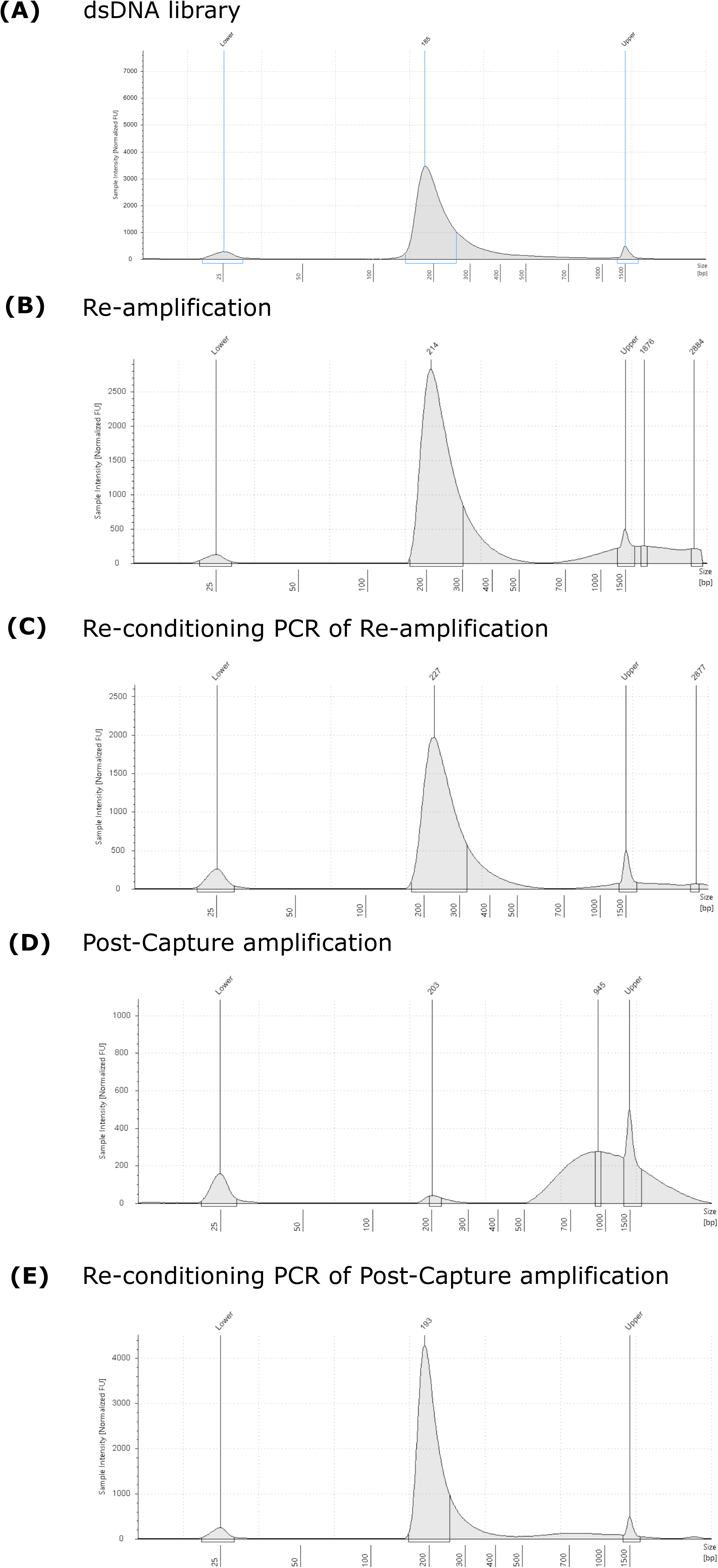
Qualitative assessment of a dsDNA library at different stages of the target enrichment process using a 4200 TapeStation instrument. **(A)** TapeStation output of the shotgun dsDNA library. **(B)** TapeStation output after the re-amplification of a shotgun dsDNA library. **(C)** TapeStation output after applying the re-conditioning PCR amplification on a re-amplified dsDNA library. **(D)** TapeStation output after the amplification of an enriched dsDNA library. **(E)** TapeStation output after applying the re-conditioning PCR amplification on an enriched dsDNA library.

## Supplementary Datafiles

**Supplementary Data SD1. Summary of laboratory information and sequencing information for each sample included in this study. (A)** Sample information and information on the extraction methods were obtained at the Institute of Genomics, University of Tartu (Estonia) and Uppsala University (Sweden). **(B)** Information on preparing dsDNA libraries using a modified version (Meyer and Kircher, 2010b) and the re-amplification concentrations processed at Uppsala University (Sweden). **(C)** Information on the target enrichment using the Twist and Tweap protocol. **(D)** Sequencing information of shotgun and enriched dsDNA libraries. **(E)** Number of SNPs and enrichment factor using the raw count. (F) Number of SNPs and enrichment factor using read normalisation. bp = base pairs. no = number. R = reamplification. SNPs = single-nucleotide polymorphisms.

